# Mitochondria biogenesis in the synapse is supported by local translation

**DOI:** 10.1101/789164

**Authors:** Bozena Kuzniewska, Dominik Cysewski, Michal Wasilewski, Paulina Sakowska, Jacek Milek, Tomasz M. Kulinski, Pawel Kozielewicz, Michal Dadlez, Agnieszka Chacinska, Andrzej Dziembowski, Magdalena Dziembowska

## Abstract

Synapses are the regions of the neuron that enable the transmission and propagation of action potentials on the cost of high energy consumption and elevated demand for mitochondrial ATP production. The rapid changes in local energetic requirements at dendritic spines imply the role of mitochondria in the maintenance of their homeostasis. Using global proteomic analysis supported with complementary experimental approaches, we show that an important pool of mitochondrial proteins is locally produced at the synapse indicating that mitochondrial biogenesis takes place locally to maintain the pool of functional mitochondria in axons and dendrites. Furthermore, we show that stimulation of synaptoneurosomes induces the local synthesis of mitochondrial proteins that are transported to the mitochondria and incorporated into the protein supercomplexes of the respiratory chain.

**Synopsis:** Mitochondria biogenesis in the synapse is supported by local translation

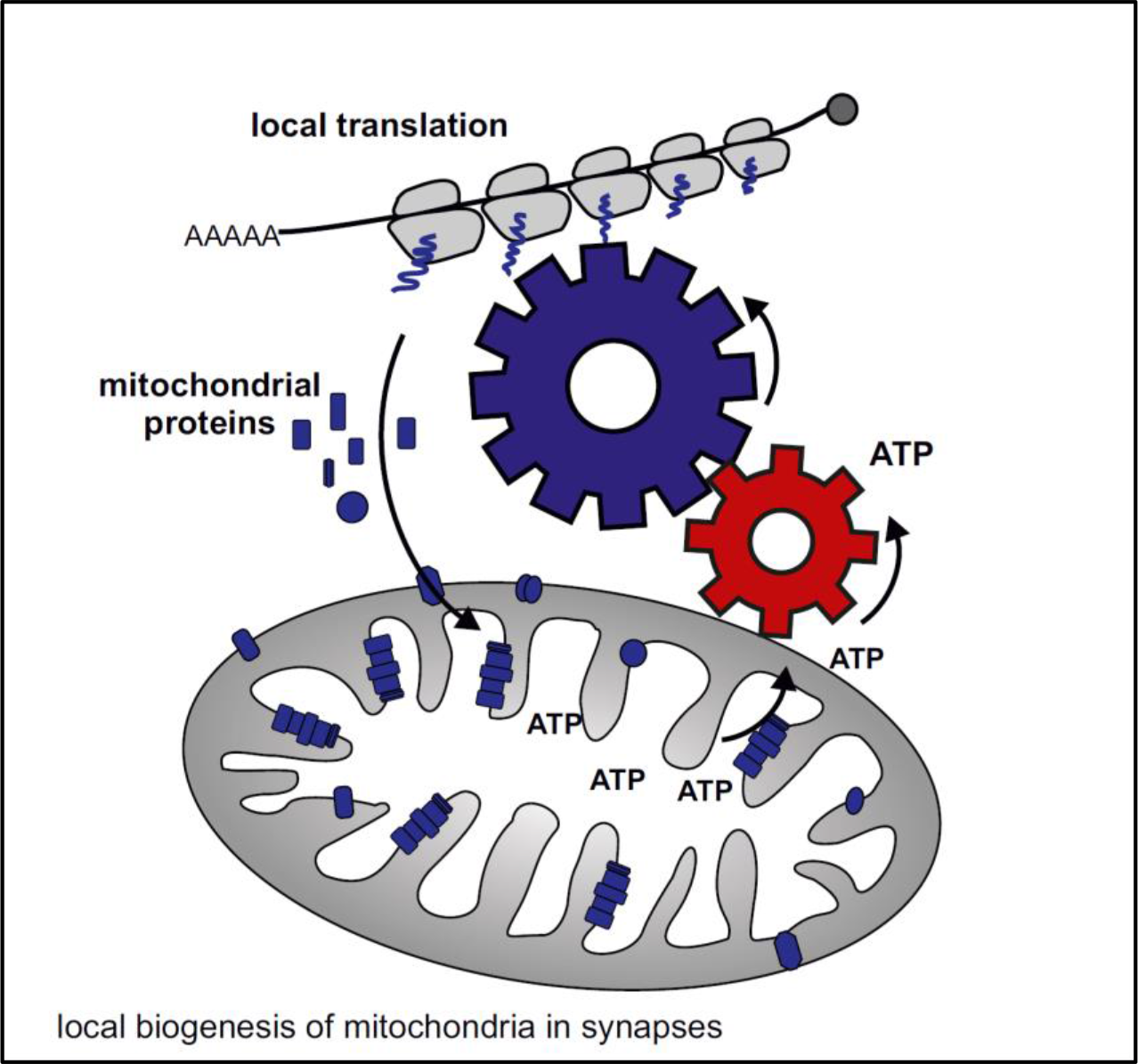

- Mitochondrial proteins represent a large pool of proteins synthesized locally at the synapse
- Newly synthesized mitochondrial proteins are imported into the mitochondria and incorporated into the respiratory chain complexes
- Uncoupling of mitochondria and blocking mitochondrial import inhibits incorporation of *de novo* synthesized proteins into the mitochondrial protein complexes

## Introduction

Synapses are spatialized zones of communication between neurons that enable the transmission and propagation of the signals. Recently it was shown, that synapses are the regions of the neuron with the highest energy consumption. Thus they have the highest demand for mitochondrial ATP production (Hollis, Kanellopoulos et al., 2017, Misgeld & Schwarz, 2017, Zhu, Qiao et al., 2012). More specifically it is the synaptic excitability that provokes temporal ion influx that will require millions of ATP molecules to be hydrolyzed to pump the ions back across the plasma membrane (Zhu et al., 2012). Maintaining resting potentials and firing action potentials are energetically expensive, as is neuro-transmission on both the pre- and postsynaptic sides (Howarth, Gleeson et al., 2012). The rapid changes in local energetic demands at dendritic spines imply the role of mitochondria in the maintenance of their homeostasis.

Synapses underlay the plastic changes phenomenon referred to as synaptic plasticity. Some forms of synaptic plasticity require mRNA translation in the postsynaptic region (Richter & Klann, 2009, Sutton & Schuman, 2006). This process proved to be extremely important for the physiology of neurons, and its dysfunction leads to abnormalities observed in the disease syndromes such as fragile X syndrome (FXS, a mutation in Fragile X mental retardation 1 gene, *FMR1*) and autism (Hagerman, Berry-Kravis et al., 2017). The discovery of actively translating polyribosomes in dendritic spines raised the question which mRNAs are locally translated and what is their function (Steward & Levy, 1982). Since then many experimental approaches have been applied to identify mRNAs transported to the dendrites and translated upon synaptic stimulation. Transcriptomic studies suggested that a large portion of proteins present in dendrites and at dendritic spines can be synthesized locally on the base of mRNA specifically transported to this compartment (Cajigas, Tushev et al., 2012). These results reveal a previously unappreciated enormous potential for the local protein synthesis machinery to supply, maintain and modify the dendritic and synaptic proteome. However, one of the major questions in this field that still remains uncovered is a global impact of local translation on synaptic functions. Recently the role of mitochondria in the plasticity of dendritic spines was revealed on the postsynapse (Rangaraju, Lauterbach et al., 2019) and in axons where the late endosomes were shown to serve as a platform for local translation (Cioni, Lin et al., 2019).

Herein, we employed quantitative mass spectrometry and *in vitro* stimulation of isolated mouse synapses (synaptoneurosomes) to create a comprehensive view of local protein synthesis in neurons. Strikingly, the second most numerous group of proteins synthesized in the synapses represented ones imported into the mitochondria. The proteomic data were further supported by sequencing of mRNAs bound with actively translating polyribosomes. Our results show that an important pool of mitochondrial proteins is locally produced at the synapse indicating that mitochondrial biogenesis takes place locally in order to maintain the pool of functional mitochondria in axons and dendrites. We further show that stimulation of synaptoneurosomes induces the local synthesis of mitochondrial proteins that are transported to the mitochondria and incorporated into of the respiratory chain complexes. This contributes to mitochondrial biogenesis in neurons and can be a potential reason why in many neurodevelopmental conditions the dysregulation of mitochondrial function is observed (Hollis et al., 2017, Khacho, Harris et al., 2019).

## Results and Discussion

### Mitochondrial proteins represent a significant fraction of locally synthesized proteins in synaptoneurosomes

To explore the dynamics of synaptic proteome upon neuronal stimulation we employed a simple model system, isolated synaptoneurosomes (SN). Synaptoneurosomes are a fraction of brain homogenate obtained by the series of filtrations and centrifugations. The resulting fraction is enriched in synapses, containing both pre- and postsynaptic compartments (Fig. 1A). The western blot on fractions obtained during SN preparation revealed the enrichment of both pre and postsynaptic markers in the SN fraction (Fig. 1B). Directly after the isolation, synaptoneurosomes can be *in vitro* stimulated to induce local protein translation. We have chosen the stimulation protocol that promotes the induction of N-methyl-D-aspartate receptors (NMDA-Rs) on the synaptoneurosomes which initiates calcium signaling in neurons and in physiological conditions, in the brain, leads to long-lasting responses such as long term potentiation (LTP) (Gustafsson & Wigstrom, 1988). For this, we treated synaptoneurosomes for 30 seconds with NMDA and glutamate and added a selective NMDA-R antagonist (APV) to avoid the induction of excitotoxicity. This treatment produces the transient phosphorylation of extracellular signal-regulated protein kinases (ERKs) in synaptoneurosomes which reflects activity-induced calcium influx mediated by NMDA receptors (Supplementary Figure S1) (Hardingham, Arnold et al., 2001, Krapivinsky, Krapivinsky et al., 2003). Next, in order to study activity-induced protein translation we incubated synaptoneurosomes with radioactive methionine/cysteine prior to NMDA-R stimulation. We observed incorporation of radioactive aminoacids into the newly synthesised proteins at 15, 30, 60 and 120 minutes as revealed by the autoradiography of the SDS-PAGE gel (Fig. 1C). In the control experiments we incubated SN with chloramphenicol, and observed no major effect of this mitochondrial protein synthesis inhibitor on the level of protein translation at 1 hour after the stimulation. However when synaptoneurosomes were incubated (pretreated) with puromycin, a significant inhibition of the translation was observed (Fig. 1C).

**Figure 1.**
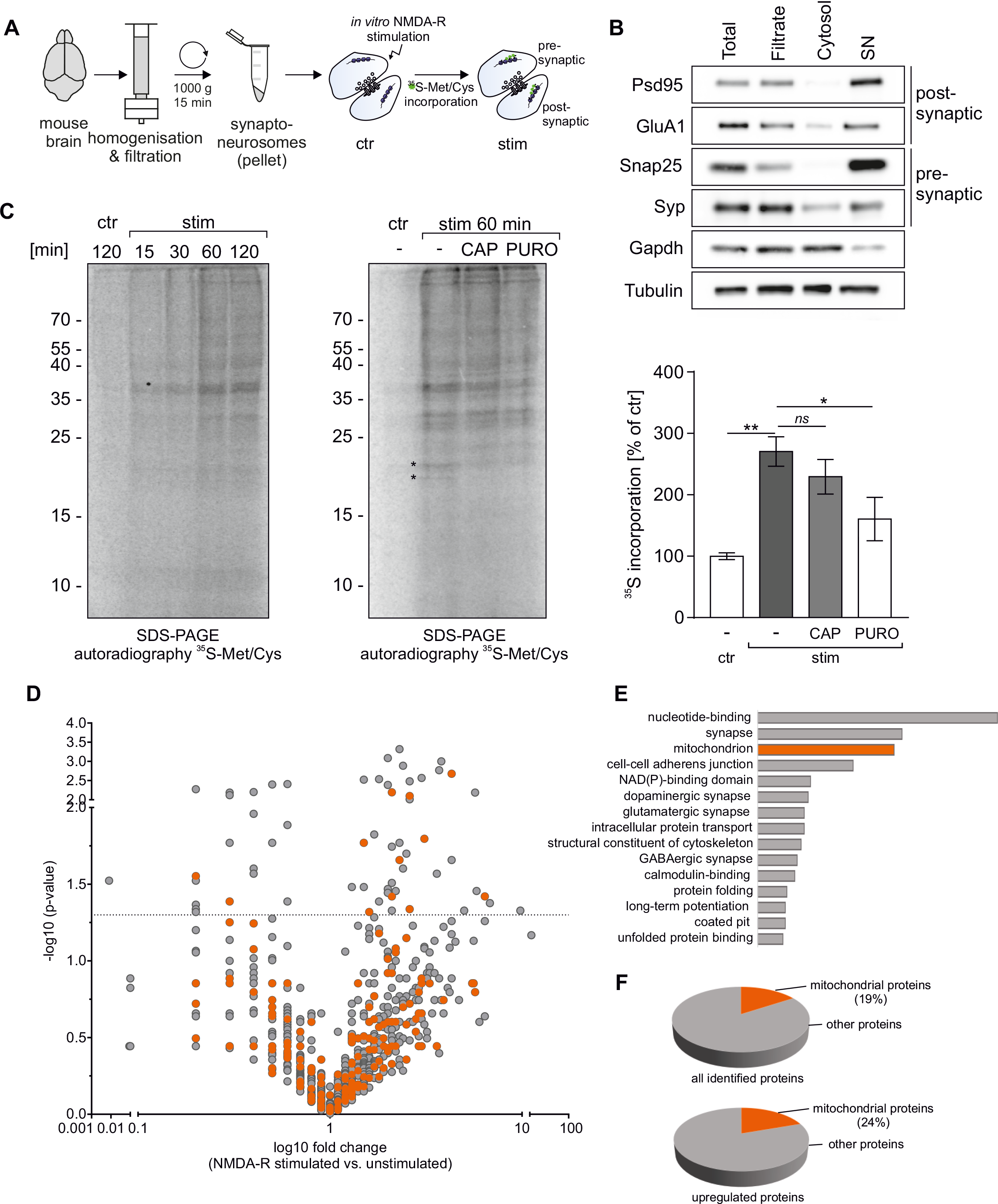
Mitochondrial proteins represent a significant fraction of locally synthesized proteins in synaptoneurosomes. (A) Workflow of the experiment presented in panel C, depicting synaptoneurosomes (SN) isolation and *in vitro* stimulation in the presence of radioactive ^35^S-labelled methionine/cysteine mix. (B) Western blot on fractions obtained during SN preparation revealed the enrichment of both pre- and postsynaptic markers in the SN. (C) Synaptoneurosomes were NMDAR-stimulated and incubated with radioactive ^35^S-methionine/cysteine mix for 15-120 min. After labeling proteins were separated on SDS-PAGE and newly synthesized proteins in SN labeled with [^35^S] were visualized by autoradiography (left panel). To determine the contribution of mitochondrial and cytosolic translation the labeling was preceded by treatment with chloramphenicol (CAP, 50 μg/ml) or with puromycin (PURO, 3mM) respectively (right panel). After 1 hour after the stimulation, significant increase in ^35^S-labelled proteins was observed (n=5, ** p<0.01, one-way ANOVA, *post-hoc* Sidak’s multiple comparisons test). CAP treatment did not affect the overal levels of *de novo* synthetized proteins (n=5, *ns* p>0.05, one-way ANOVA, *post-hoc* Sidak’s multiple comparisons test). Please note however, that there are two bands missing in CAP-treated SN samples (marked by asterisks) as compared to the stimulated sample. In contrast, PURO treatment significantly inhibited *de novo* protein synthesis in synaptoneurosomes (n=5, * p<0.05, one-way ANOVA, *post-hoc* Sidak’s multiple comparisons test). (D-F) Results of mass spectrometry (label free quantification - LFQ) analysis identifying proteins with increased abundance in stimulated NMDAR-stimulated synaptoneurosomes. (D) Volcano plot showing abundance of identified proteins in stimulated synaptoneurosomes as compared to unstimulated. Mitochondrial proteins are depicted in orange. The vertical line defines the p-value statistical significance cut-off. (E) Biological functions of identified proteins annotated using DAVID. Third top functional category identified as mitochondrial ones. (F) 19% of all identified proteins were mitochondrial and 24% of all upregulated ones.

Protein translation in isolated synaptoneurosomes depleted of nuclei and cell bodies rely on the pool of mRNAs transported into the synapses in the brain, therefore we reasoned that all proteins which abundance will increase after the *in vitro* stimulation would be locally synthesized on the base of these mRNAs. Thus, in order to identify the proteins newly synthetized in stimulated synaptoneurosomes we performed mass spectrometry (label free quantification - LFQ) and compared the abundance of proteins in unstimulated and NMDA-R stimulated SN (Fig. 1D). Total number of identified proteins was 2200 (FDR 1%). In order to gain some insight in the biological functions of identified proteins, we have performed standard gene annotation using DAVID. Strikingly the third most numerous group was annotated as proteins imported to the mitochondria, after the nucleotide-binding and synaptic proteins (Fig.1E). 19% of all identified proteins were mitochondrial and 24% of all upregulated ones (Fig.1F).

Because of the small number of proteins that reached statistical significance in the LFQ analysis we turned into the quantitative mass spectrometry based on isobaric labeling techniques and we performed labeling of the samples with iTRAQ or TMT tag (Figure 2A and B). Quantitative mass spectrometry heavily depends on physicochemical properties of peptides and dynamic range of samples which limits in the depth of analysis. Some of tryptic peptides would be better detectable unmodified, other when modified for example with stable isobaric labeling techniques such as iTRAQ or TMT tag. Therefore in order to gain a deeper insight into the nature of synaptoneurosomal proteome and overcome physicochemical limitations we measured the same samples using two quantitative methods: iTRAQ8 and TMT10 on the same biological material. 1942 proteins were identified using iTRAQ8-plex and 2508 in TMT10-plex. All together in three independent proteomic approaches we identified 3201 proteins out of which we were able to quantify 2080. 516 of those proteins were significantly upregulated in the stimulated synaptoneurosomal samples (Mann-Whitney p<0.05). The list of identified proteins is attaches as a supplementary Table1. Next, we performed gene-annotation enrichment analysis with DAVID on proteins identified as significantly upregulated after NMDA-R stimulation (assembled from all three proteomic analyzes). Here we observed mitochondrial proteins to be the second most abundant category of identified proteins (Fig. 2C).

**Figure 2.**
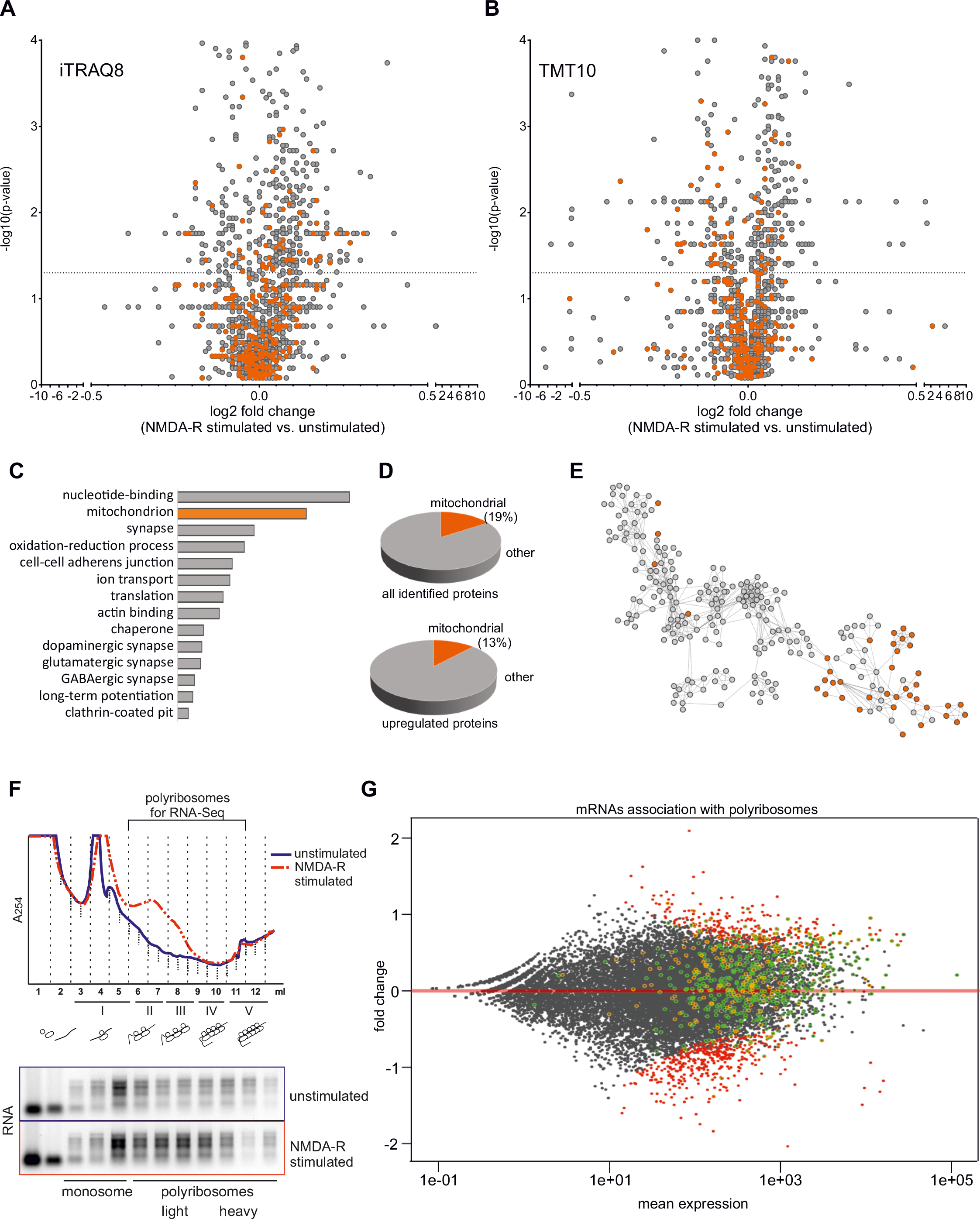
Identification of proteins with increased abundance in stimulated synaptoneurosomes using high-resolution quantitative mass spectrometry. (A-E) Results of high-resolution quantitative mass spectrometry analysis identifying proteins with increased abundance in stimulated synaptoneurosomes. Volcano plots showing abundance of identified proteins in stimulated synaptoneurosomes as compared to unstimulated using iTRAQ8-plex (A) and TMT10-plex (B) labelling methods. Mitochondrial proteins are depicted in orange. The vertical line defines the p-value statistical significance cut-off. For nine mitochondrial proteins their mRNA abundance on polyribosomes was quantified, the data is presented on Figure S2. (C) Biological functions of proteins identified as significantly upregulated after NMDA-R stimulation (assembled from all three proteomic analyzes: LFQ, iTRAQ8, TMT10) annotated using DAVID. Second top functional category identified as mitochondrial ones. (D) Mitochondrial proteins constituted 19% (393) of all quantified proteins and 13.4% (69) of upregulated ones. (E) Interactome of identified proteins, mitochondrial proteins shown in orange. (F) RNA absorbance profiles from polyribosomal fractionation of unstimulated and NMDAR-stimulated synaptoneurosomes. Upon NMDAR stimulation, fraction of mRNAs is shifted towards heavy-polyribosomal fraction, that represent actively translating ribosomes. Fraction of polyribosomes used for RNA-seq experiment is depicted. (G) Comparison of proteomic data with RNA-seq experiments on mRNAs associated with polyribosomes from synaptoneurosomal samples. Ribosome association of synaptoneurosomal mRNAs determined by normalizing RNA-seq values of polyribosome associated mRNA with total mRNA (y axis). Green dots represent mRNAs encoding proteins upregulated upon stimulation (identified in MS experimets) while yellow dots represents mRNAs encoding mitochondrial proteins. Please notice that both classes represent high abundant mRNAs with good ribosome association values. Transcripts with statistically significant increased or decreased abundance on the polyribosomes in response to the stimulation are depicted in red. *See also Figure S2.*

To look more specifically on mitochondrial proteins we equated our results to a mitochondrial database of MitoCarta 2.0. Mitochondrial proteins constituted 19% (393) of all quantified proteins and 13.4% (69) of upregulated ones (Fig.2D). This finding is remarkable since mitochondrial proteins (1160 gene names, MitoCarta 2.0) represent only 5% of all protein sequences in the database downloaded from UniProt (52539 entries) and used for the results annotation.

Proteomic mass spectrometry data from Supplementary Table 1 and the software program Cytoscape (version 3.5.1) were used to generate a graphic representation of known protein-protein interactions for proteins significantly upregulated by NMDA-R stimulation (Fig 2E).

The number of quantified proteins (3201) could be considered as relatively low for shotgun MS experiment, however one should take into account the specificity of the synaptoneurosomal preparations depleted of nuclear and probably many cytoplasmic proteins. Importantly, 516 proteins upregulated in synaptoneurosomes by the stimulation corresponds to as much as 20% of all identified proteins. The average fold change was relatively low, which was however expected because of the translational capacity of synaptic ribosomes.

To strengthen the proteomic data we additionally analyzed synaptoneurosomal transcriptome engaged into translation. For this, we took advantage of the fact that local synthesis of synaptic proteins is performed on site with the pool of available ribosomes (Steward & Fass, 1983) that can be isolated from synaptoneurosomes and fractionated on the sucrose gradient according to their molecular weight (Kuzniewska, Chojnacka et al., 2018). Thus we fractionated polyribosomes from stimulated synaptoneurosomes to identify the mRNAs associated with polyribosomal fractions in response to the stimulation (Fig. 2F). Then we performed RNA-seq experiments and calculated the increase of given mRNA in the polyribosomal fraction in response to NMDA-R stimulation comparing to total mRNA from the unstimulated sample (Fig. 2G). We detected 17160 different transcripts among which 3078 (18%) were detected at the protein level. 9312 of mRNAs were enriched in the polyribosomal fraction when compared to the total SN fractions. 64% of all upregulated proteins were also enriched on mRNA level what was an additional proof for their activity-induced polyribosomal binding and translation. In case of the remaining 36% of proteins we did not detect their enrichment in the polyribosomal fraction. Nevertheless, they were mostly already present in the polysomal fraction, probably undergoing translation on the basal level and not engaging into the polysomal fraction after the stimulation. Importantly, 56 of 70 upregulated mitochondrial proteins were also overrepresented at the mRNA level in the polyribosomal fraction (Fig. 2G). Next, we verified RNA-seq data using qRT-PCR on the polysomal fractions isolated from synaptoneurosomes (selected mRNAs are listed in Supplementary Figure 2). Indeed, we could see the transcripts encoding for mitochondrial proteins shift towards the polyribosomal fractions upon the NMDA-R stimulation (Supplementary Figure 2).

### Locally synthesized proteins build mitochondria in the synapse

The large fraction of locally synthesized mitochondrial proteins identified in our mass spectrometry analysis were the constituents of the respiratory chain complexes (see Supplementary Table 1). Mitochondria in presynaptic terminals of synapses were for a long time used as a hallmark of synapses and their real role in neurotransmission is recently being determined (Hollenbeck, 2005). The studies of axonal local translation in squid giant axon and presynaptic nerve terminals of the photoreceptor neurons revealed the presence of mRNAs population containing nuclear-encoded mitochondrial proteins (Gioio, Eyman et al., 2001, Kaplan, Gioio et al., 2009). Synapses in the brain contain mitochondria, therefore we checked the presence of the mitochondrial proteins in SN and all fractions obtained during the preparation procedure. Proteins representing respiratory chain complexes (Ndufa9, Sdha, Uqcrc1, Atp5b) and the outer membrane transporter (Tom20) were present in SN (Fig. 3A). To confirm that the mitochondria located in SN are functional we measured O2 consumption following supplementation in respiratory substrates, uncouplers and inhibitors. Quantitative respirometry was performed in parallel to compare the respiration rates between permeablized HEK293T cells and SN. We detected similar responses to the addition of respiratory substrates for complex I (malate and pyruvate) and complex II (succinate) in the case of HEK293T cells and SN pointing at their similar respiration status (Fig. 3B). We also compared the native mitochondrial protein complexes from SN and HEK cells obtained by digitonin extraction and blue-native gel electrophoresis (BN-PAGE), the golden standard for visualization of respiratory complexes and supercomplexes present in mitochondria(Figure 3C). Characteristic pattern of bands representing the native mitochondrial respiratory chain complexes appeared to be similar in SN and HEK293T cells although more high molecular weight bands in the region of supercomplexes was revealed in SN. Next, the BN-PAGE gels were electrotransfered on the PVDF membrane for western blot analysis. We used the panel of antibodies specifically recognizing the protein components of complex I (Ndufb8), complex II (Sdha), complex III (Uqcrc2), complex IV (Cox6a) and complex V (Atp5a) that revealed the presence of these complexes in the BN-PAGE gels (Fig. 3D). The pattern and identity of proteins were in agreement with a well-established profile of the native respiratory chain complexes from mouse heart (Mourier, Matic et al., 2014). Interestingly we observed more bands corresponding to supercomplexes in SN than in HEK, which confirmed the Coomassie staining (Fig. 3C and D).

**Figure 3.**
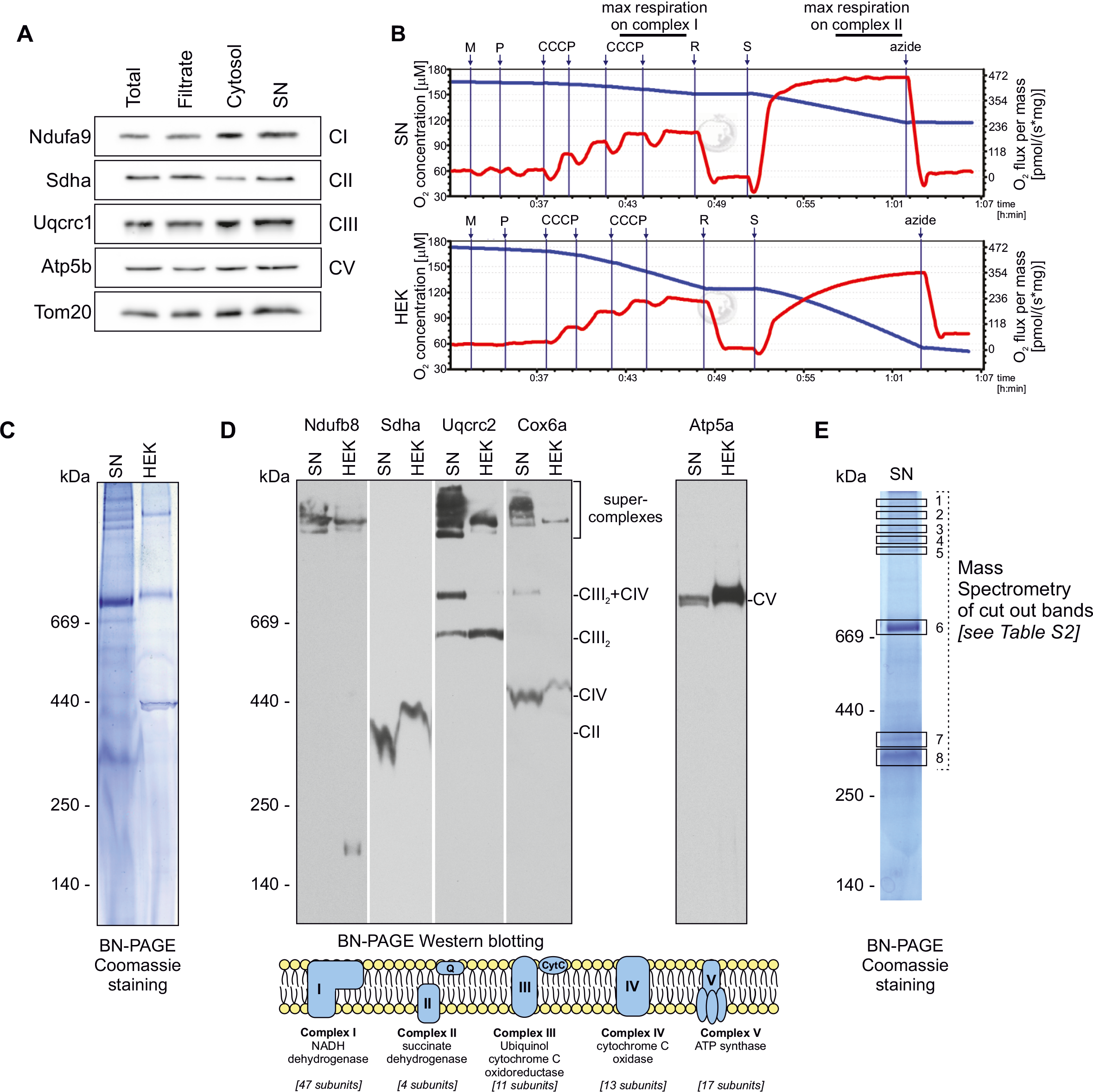
Characterization of mitochondrial respiratory complexes and their activity in synaptoneurosomes. (A) Western blot on fractions obtained during SN preparation revealed presence of respiratory chain complex components (Ndufa9, Sdha, Uqcrc1, Atp5b) and the outer membrane transporter (Tom20) in SN. (B) Respiration activity of permeablized synaptoneurosomes (upper panel) as compared to permeablized HEK293T (lower panel). Oxygen concentration (blue line) and oxygen consumption (red line) measured using O2k oxygraph are shown. (C) Blue native gel electrophoresis (BN-PAGE) analysis of mitochondrial respiratory chain complexes from synaptoneurosomes as compared to HEK293T mitochondria. (D) Detection of respiratory chain complexes and supercomplexes in SN and HEK293T mitochondria using BN-PAGE followed by Western blot using antibodies specific for: complex I (Ndufb8), complex II (Sdha), complex III (Uqcrc2), complex IV (Cox6a) and complex V (Atp5a). Note similar composition of respiratory supercomplexes and complexes in synaptoneurosomes as in cellular mitochondria. (E) Selected bands (1-8) visible in blue-native gel were excised and analyzed using mass spectrometry (LC-MS). Proteomic analysis confirms the presence of protein subunits of different respiratory chain complexes in indicated bands. *See Table S2 for identified peptides.*

For the undeniable characterization of the protein complexes visible on the BN-PAGE gels we dissected eight bands from the polyacrylamide gel and analyzed them by mass spectroscopy (Fig 3E). The protein components of mitochondrial respiratory chain complexes were identified (listed in the Supplementary Table 2).

To establish that locally translated mitochondrial proteins are effectively transported into the mitochondria we loaded synaptoneurosomes with radioactive methionine/cysteine, stimulated NMDA-Rs allowing local protein synthesis, and assayed for incorporation of radioactive proteins into the native mitochondrial respiratory chain complexes on BN-PAGE (Figure 4A). The incorporation of newly synthesized radiolabeled proteins into complexes detected by BN-PAGE resulted in a pattern overlapping with the mature complexes detected by the Coomassie staining (Fig. 3C). Additional bands formed by radiolabeled proteins likely represented import and complex assembly intermediates (Mick, Fox et al., 2011). The respiratory complexes consist of proteins that are also encoded by mitochondrial DNA. As the control, the SN were treated with CCCP or VOA to abolish mitochondrial electrochemical potential and thus inhibit protein transport into the mitochondria. This treatment effectively inhibited the activity-induced appearance of radioactive respiratory chain complexes containing newly synthesised proteins in the mitochondria (Fig. 4B and C). Altogether, we established that synaptically translated proteins are imported into mitochondria and assembled into the functional respiratory chain complexes.

**Figure 4.**
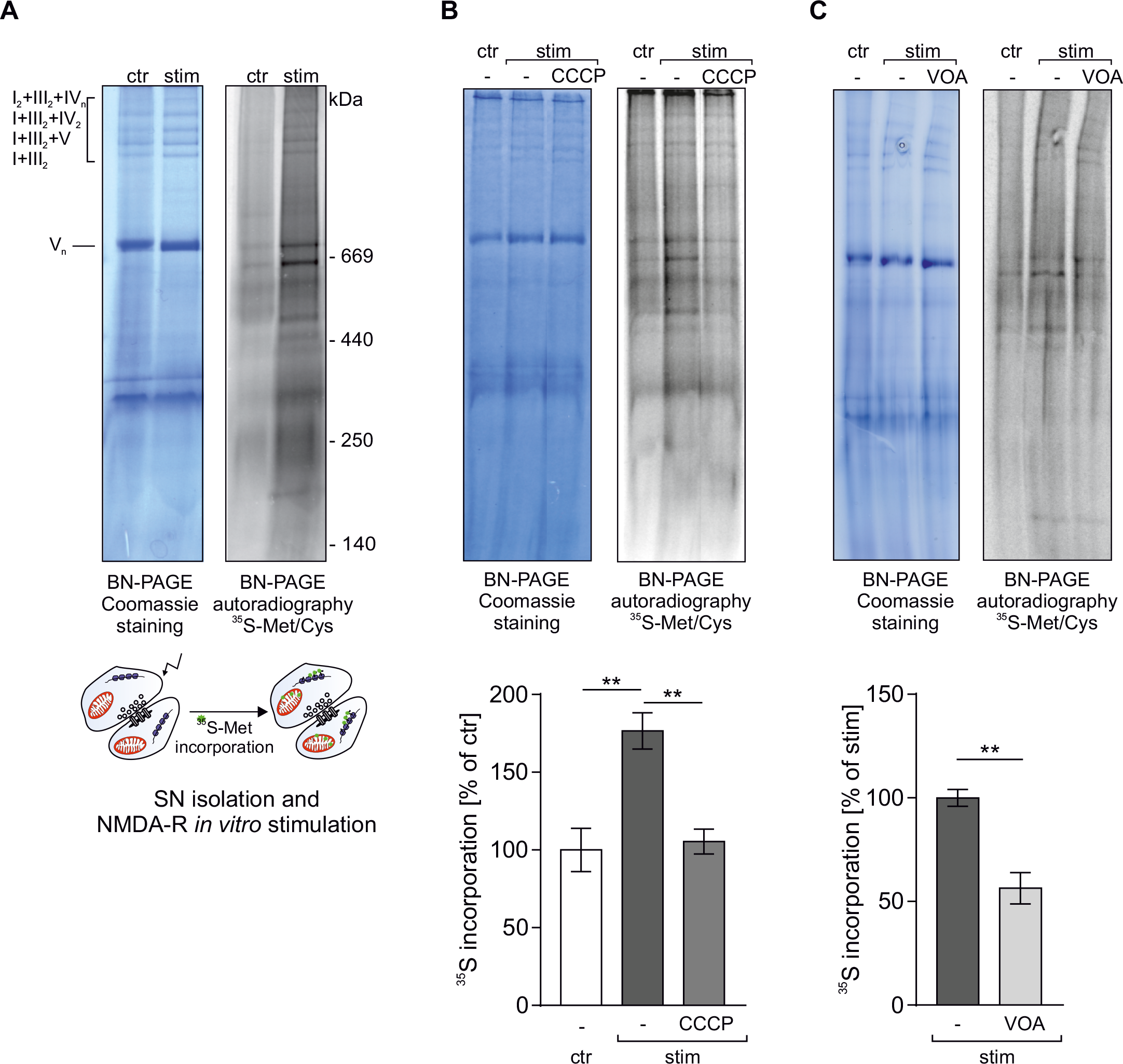
Mitochondrial proteins locally synthesized in the synapse contribute to mitochondrial biogenesis. (A) Synaptoneurosomes were NMDAR-stimulated and incubated with radioactive ^35^S-methionine/cysteine mix for 1h. Samples were separated using blue-native electrophoresis (BN-PAGE). Protein complexes were visualized by Coomassie staining (left) and autoradiography (right). Autoradiography of the BN-PAGE gel shows *de novo* synthetized mitochondrial proteins that are incorporated into respiratory chain complexes. (B-C) To block protein import and incorporation of newly synthesized proteins into respiratory complexes the mitochondrial electrochemical potential was abolished by treatment with carbonyl cyanide m-chlorophenyl hydrazone (10 μM CCCP) (B) or VOA mixture (containing 1 μM valinomycin, 20 μM oligomycin, 8 μM antimycin) (C). Protein complexes were separated on BN-PAGE and visualized by Coomassie staining and autoradiography. Significant inhibition of ^35^S-methionine/cysteine incorporation into mitochondrial protein complexes in synaptoneurosomes was observed when samples were incubated in the presence of CCCP (B; n=4, ** p<0.01, One-way ANOVA, *post-hoc* Sidak’s multiple comparisons test) or VOA (C; n=3, ** p<0.01, One-way ANOVA, *post-hoc* Sidak’s multiple comparisons test).

## Conclusions

In this study, we have made a serendipitous discovery of mitochondria biogenesis at the neuronal synapse. Interestingly, mitochondrial proteins not only represent one of the most abundant class of locally synthesized proteins in response to neuronal stimulation but they are also capable of entering mitochondria followed by the effective formation of functional complexes (graphical abstract). In addition, our study defines mitochondrial biogenesis as a new group of potential therapeutic targets in diseases such as fragile X syndrome where the synaptic translation is dysregulated.

## Materials and methods

### Animals

1- to 2-month-old male FVB mice (FVB/NJ, Jackson Laboratories Stock No.: 001800) were used. Before the experiment, the animals were kept in the laboratory animal facility under a 12-h light/dark cycle with food and water available *ad libitum*. The animals were treated in accordance with the EU Directive 2010/63/EU for animal experiments.

### Preparation of synaptoneurosomes and stimulation of NMDA receptors

Synaptoneurosomes were prepared as described previously (Dziembowska, Milek et al., 2012, Kuzniewska et al., 2018, Scheetz, Nairn et al., 2000). Before tissue dissection Krebs buffer (2.5 mM CaCl_2_, 1.18 mM KH_2_PO_4_, 118.5 mM NaCl, 24.9 mM NaHCO_3_, 1.18 mM MgSO_4_, 3.8 mM MgCl_2_, 212.7 mM glucose) was aerated with an aquarium pump for 30 min at 4°C. Next, the pH was lowered to 7.4 using dry ice. The buffer was supplemented with 1×protease inhibitor cocktail cOmplete EDTA-free (Roche) and 60 U/ml RNase Inhibitor (RiboLock, Thermo Fisher Scientific). Animals were euthanized by cervical dislocation, hippocampi and a part of cortex adjacent to the hippocampus were dissected. Tissue from one hemisphere (~50 mg) was homogenized in 1.5 ml Krebs buffer using Dounce homogenizer with 10-12 strokes. All steps were kept ice-cold to prevent stimulation of synaptoneurosomes. Homogenates were loaded into 20 ml syringe and passed through a series of pre-soaked (with Krebs buffer) nylon mesh filters consecutively 100, 60, 30 and 10 μm (Merck Millipore) in cold room to 50 ml polypropylene tube, centrifuged at 1000 g for 15 min at 4°C, washed and pellet was resuspended in Krebs buffer with protease and RNase inhibitors. The protocol for *in vitro* stimulation of NMDA receptors on synaptoneurosomes was described before (Kuzniewska et al., 2018, Scheetz et al., 2000). The aliquots of freshly isolated synaptoneurosomes were prewarmed for 5 min at 37°C and stimulated with a pulse of 50 μM NMDA and 10 μM glutamate for 30 sec, then APV (120 μM) was added and synaptoneurosomes were further incubated for indicated time at 37°C. Unstimulated samples, kept on ice with 200 μg/ml cycloheximide, were used as controls.

### Western blot analysis of synaptoneurosome preparations

Equal amounts of protein from homogenate, filtrate, supernatant (cytosol fraction) and synaptoneurosomal fraction were resolved on SDS-PAGE (10%, TGX Stain-Free FastCast Acrylamide Solutions, BioRad). After electrophoresis proteins in the gel were visualized using Bio-Rad’s ImageLab software to verify the equal protein loading. Proteins were transferred to PVDF membranes (pore size 0.45 μm, Immobilon-P, Merck Millipore) using Trans-Blot Turbo Blotting System (BioRad; 170-4155). Membranes were blocked for 1 h at room temperature in 5% non-fat dry milk in PBS-T (PBS with 0.01% Tween-20), followed by overnight incubation at 4°C with primary antibodies (PSD95, Snap25, synaptophysin, Gapdh, tubulin, Ndufa9, Sdha, Uqcrc1, Atp5b, Tom20) in 5% milk in PBS-T. Blots were washed 3 × 5 min with PBS-T, incubated 1 h at room temperature with HRP-conjugated secondary antibody (1:10 000 in 5% milk) and washed 3 × 5 min with PBS-T. In the case of MAPK/ Phospho-MAPK Family Antibody Sampler Kit and anti-GluA1 antibody, membranes were blocked for 1 h at room temperature in 5% bovine serum albumin (BSA) in PBS-T, primary and secondary antibodies were diluted also in 5% BSA in PBS-T. HRP signal was detected using Amersham ECL Prime Western Blotting Detection Reagent (GE Healthcare) on Amersham Imager 600 using automatic detection settings.

### Protein labeling with ^35^S-Methionine/Cysteine and autoradiography

Freshly isolated synaptoneurosomes were incubated with 100 μCi [^35^S] methionine/cysteine mix (Perkin Elmer) for 10 minutes at 4°C. Next, synaptoneurosomes were prewarmed for 5 minutes at 37°C and stimulated (50 μM NMDA and 10 μM glutamate for 30 sec, followed by addition of 120 μM APV). Synaptoneurosomes were further incubated at 37°C for indicated time points. A control sample was stimulated and kept on ice. Synaptoneurosomes were subsequently centrifuged (1000 × g, 10 min at 4°C), washed with ice-cold Krebs buffer with protease and RNase inhibitors, centrifuged again and further subjected to SDS-PAGE or BN-PAGE followed by autoradiography or western blot with specific antibodies.

Where indicated, isolated synaptoneurosomes were stimulated in the presence of 50 μg/ml chloramphenicol (CAP), 3mM puromycin (PURO), 10 μM carbonyl cyanide m-chlorophenyl hydrazone (CCCP) or VOA mixture (containing 1 μM valinomycin, 20 μM oligomycin, 8 μM antimycin).

For SDS-PAGE proteins were precipitated with pyrogallol red-molybdate (0.05 mM pyrogallol red, 0.16 mM sodium molybdate, 1 mM sodium oxalate, 50 mM succinic acid) for 20 minutes and room temperature, centrifuged (20000 × g for 15 minutes), resuspended in urea sample buffer (6M urea, 6% SDS, 125 mM Tris-HCl [pH 6.8], 0.01% bromophenol blue, 50 mM DTT) and separated on 15% polyacrylamide gels. Separated proteins were fixed using Coomassie staining solution. Gels were further destained, dried in a vacuum dryer and incubated with a Storage Phosphor Screen (GE Healthcare). Radioactively labeled proteins were visualized by digital autoradiography (Storm imaging system; GE Healthcare) followed by image processing with ImageQuant software (GE Healthcare).

For analysis of native protein complexes (blue native electrophoresis, BN-PAGE) synaptoneurosomes and isolated mitochondria were resuspended in solubilization buffer (5% [wt/vol] digitonin, 20 mM Tris-HCl, pH 7.4, 50 mM NaCl, 10% [wt/vol] glycerol, 0.5 mM EDTA, 2 mM PMSF) and incubated for 20 minutes on ice. Mitochondria from HEK293T cells were isolated as described in (Mohanraj, Wasilewski et al., 2019).

Samples were centrifuged at 12,000 × g at 4°C and the soluble fraction was mixed with loading dye (5% [w/v] Coomassie Brilliant Blue G-250, 100 mM Bis-Tris, and 500 mM ε-amino-n-caproic acid [pH 7.0]). HMW Native Marker Kit (GE Healthcare) was used for protein complexes weight reference. Samples were separated on a 4–13% gradient gel at 4°C. For LC-MS analysis, separated protein complexes were subjected to Coomassie staining. After destaining, the bands representing protein complexes were excised from a gel and subjected to LC-MS. Radioactively labeled proteins were visualized by autoradiography as described for SDS-PAGE gels. For western blotting, protein complexes were transferred to polyvinylidene difluoride membranes (PVDF, Millipore) by a semi-dry transfer (250 mA for 2 hours) in the Blotting buffer (20mM TRIS; 0.15 M glycine, 10% methanol). Membranes were incubated with specific primary antibodies and detected by an enhanced chemiluminescence detection system using X-ray films (Foton-Bis). Custom rabbit polyclonal antibodies (from Peter Rehling Lab): anti-COX6A (1:2000; PRAB3283), anti-NDUFB8 (1:1000; PRAB3765), and commercial mouse monoclonal antibody: ATP5A (15H4C4, Abcam, 1:500), SDHA (D-4, Santa Cruz, 1:2000), UQCRC2 (13G12AF12BB11, Abcam, 1:500) were used.

### Proteomics

Synaptoneurosomes were frozen at −80°C, 500 μl of buffer (25 mM HEPES, 2% SDS, protease and phosphatase inhibitors) was added. Samples were heated at 96°C for 3 minutes, cooled down and sonicated in Bioruptor® Plus (Diagenode) for 20 cycles 30/30 seconds at “high”; then samples were again heated for 3 minutes at 96°C.

Protein concentration has been determined using DirectDetect Spectrometer (Merck Millipore). Appropriate volumes containing accordingly 75 μg proteins and 10 μg, respectively for isobaric labeling and label-free experiments, per sample were moved to 1.5 ml tubes and precipitated using chloroform/methanol protocol. Samples then were label using standard iTRAQ 8-plex (SCIEX) and TMT 10-plex (ThermoFisher) protocol according to the manufacturer’s recommendations. Label-free samples were dissolved in 100 μl of 100 mM ammonium bicarbonate buffer, reduced in 100 mM DTT for 30 min at 57°C, alkylated in 55 mM iodoacetamide for 40 min at RT in the dark and digested overnight with 10 ng/ml trypsin (V5280, Promega) at 37°C. Finally, to stop digestion trifluoroacetic acid was added at a final concentration of 0.1%. Mixture was centrifuged at 4°C, 14 000 g for 20 min, to remove solid remaining.

Samples labeled with TMT and iTRAQ were fractionated prior to LC-MS using strong cation-exchange chromatography (SCX). Briefly, 200 μg of samples were injected to PolyLC columns (2.1 × 4.6 × 200 mm, 5 μm, 300 Å pore size) and were fractionated in 60 minutes linear gradient, from 100% A (5 mM KH_2_P0_4_/ 25% acetonitrile, pH 2.8 or 0.1% formic/30% acetonitrile) to 100% B (5 mM KH_2_P0_4_/ 25% acetonitrile, pH 2.8 + 350mM KCl or 500 mM ammonium formate/ 30% acetonitrile).

Proteins in bands excised from a gel were reduced with 100 mM DTT (for 30 minutes at 57°C), alkylated with 0.5 M iodoacetamide (45 min in a dark at room temperature) and digested overnight in 37°C with 10 ng/μl of trypsin in 25mM NH_4_HCO_3_ (sequencing Grade Modified Trypsin Promega V5111) by adding the enzyme directly to the reaction mixture. Peptides were eluted with 2% acetonitrile in the presence of 0.1% TFA. The resulting peptide mixtures were applied to RP-18 pre-column (Waters, Milford, MA) using water containing 0.1% TFA as a mobile phase and then transferred to a nano-HPLC RP-18 column (internal diameter 75 μM, Waters, Milford MA) using ACN gradient (0 – 35% ACN in 180 min) in the presence of 0.1% FA at a flow rate of 250 nl/min. The column outlet was coupled directly to the ion source of Orbitrap Velos mass spectrometer (Thermo Electron Corp., San Jose, CA) working in the regime of data-dependent MS to MS/MS switch and data were acquired in the m/z range of 300–2000.

MS analysis was performed by LC-MS in the Laboratory of Mass Spectrometry (IBB PAS, Warsaw) using a nanoAcquity UPLC system (Waters) coupled to an QExactive Orbitrap mass spectrometer (Thermo Fisher Scientific). The mass spectrometer was operated in the data-dependent MS2 mode, and data were acquired in the m/z range of 100-2000. Peptides were separated by a 180 min linear gradient of 95% solution A (0.1% formic acid in water) to 45% solution B (acetonitrile and 0.1% formic acid). The measurement of each sample was preceded by three washing runs to avoid cross-contamination. Data were analyzed with the Max-Quant (Versions 1.5.6.5 and 1.5.7.4) platform using mode match between runs (Cox & Mann, 2008). The mouse proteome database from UniProt was used (downloaded at 2017.01.20). Modifications were set for methionine oxidation, carbamidomethyl or methylthio on cysteines, phospho (STY). Label-Free-Quantification (LFQ) intensity values were calculated using the MaxLFQ algorithm (Cox, Hein et al., 2014). Samples labeled with TMT or iTRAQ were searched against the same database with a 0.001Da error on isobaric tag. Results (LFQ, iTRAQ, TMT) were analyzed using Scaffold 4 platform (Proteome Software) and merge in a one session file.

### High resolution respirometry

O_2_ consumption rates in synaptoneurosomes were measured polarographically using a high-resolution respirometer (Oroboros Oxygraph-O2K). Freshly isolated synaptoneurosomes (~0.4 mg/ml) were pelleted at 1000 g for 10 min at 4°C and resuspended in mitochondria respiration medium MIR05 (0.5 mM EGTA, 3 mM MgCl_2_, 60 mM lactobionic acid, 20 mM taurin, 10 mM KH_2_PO_4_, 110 mM sucrose, BSA 0.1%, 20 mM HEPES/KOH, pH 7.1). The amount of synaptoneurosomes was approximated by OD at 600 nm and the volume corresponding to 150 μg per respirator chamber was loaded. Synaptoneurosomes were permeabilized in the chamber by digitonin (0.005%, wt/vol) for 20 min. HEK293T cells were harvested by trypsinization, counted and resuspended in the MIR05 medium. 2 mln of cells were loaded per respirator chamber and permeablized by saponin (15 μg/ml) for 20 min. Respiration was measured at 37°C. Malate (0.5 mM), pyruvate (10 mM) were added to the respirator chambers to support electron entry through complex I followed by titration of CCCP to a final concentration of 2 μM. After inhibition of complex I by rotenone (0.5 μM) succinate (10 mM) was added to support electron entry through complex II. Measurement was concluded with sodium azide (50 mM) to inhibit complex IV. Data recording was performed using Oxygrap-2k and analyzed with DatLab 7 software (Oroboros Instruments). Protein content of synaptoneurosomal and HEK293T samples was measured using Pierce BCA protein assay kit.

### Polyribosome profiling

Polyribosomal profiling of synaptoneurosomes was performed as described previously [Kuzniewska et al., 2018]. Synaptoneurosomal samples (unstimulated or NMDA-R stimulated) were centrifuged at 1000 g for 15 min at 4°C and the pellet was lysed in 1 ml of lysis buffer [20 mM Tris-HCl pH 7.4, 2 mM DTT, 125 mM NaCl, 10 mM MgCl_2_, 200 μg/ml cycloheximide, 120 U/ml RNase inhibitor, 1×protease inhibitor cocktail and 1.5% IGEPAL CA-630 (Sigma Aldrich)]. After centrifugation at 20 000 g for 15 min at 4°C, the supernatant was collected. For RNA-Seq experiment, 0.25 vol. of the supernatant was retained as “total” fraction and the remaining 0.75 vol. was loaded on top of a 10–50% linear density sucrose gradient and was centrifuged at 38 000 rpm for 2 h at 4°C in an SW41 rotor (Beckman Coulter). In the case of polyribosomal profiling using qRT-PCR whole supernatant was loaded on the gradient. After ultracentrifugation, each gradient was separated and collected into 24 fractions (0.5 ml each) using the BR-188 Density Gradient Fractionation System (Brandel) set up using following parameters: pump speed, 1.5 ml/min; chart speed, 150 cm/h; sensitivity 0.2; peak separator off; noise filter, 1.5. Simultaneously, a continuous absorbance profile of each gradient at 254 nm was graphed. Basing on this profile, fractions were combined into five pools (depicted on Figure 2): (I) monosome, (II) and (III) representing light polysomes, (IV) and (V) corresponding to heavy polysomes, that are engaged in active translation. Obtained fractions were used for RNA isolation and qRT-PCR analysis. For RNA-Seq experiment light and heavy polyribosomal fractions without RNA granules (II, III, IV and first 1 ml of fraction V) were used (depicted in Figure 2).

### RNA isolation, library preparation and RNA-sequencing

Synaptoneurosomes isolated from cortices and hippocampi of 6 mice were divided into 4 aliquots containing an equal amount of material. One aliquot was saved as a “total” synaptoneurosomal fraction from which total RNA was extracted with phenol/chloroform mixture and another was NMDA-R stimulated for 20 min. Triplicates of ribo-depleted total RNA isolated from synaptoneurosomes (“total”) as well as RNA from the NMDA-R stimulated synaptoneurosome polisome fractions were used to prepare strand-specific libraries (dUTP RNA) (Szczesny, Kowalska et al., 2018).

Polysomal fractions were isolated as described above, pooled and precipitated with 3 volumes of ethanol. Precipitates were digested with proteinase K and total RNA was extracted with phenol/chloroform mixture. Phase separation was performed using Phase Lock Gel Heavy 2 ml Tubes (5Prime). DNA contamination from 2 μg of nucleic acids was removed by 2 U of TURBO DNase (AM2238, Ambion) in 20 μl of the supplied buffer in 37°C for 30 min. RNA was extracted with phenol/chloroform, precipitated with ethanol and resuspended in RNase free water. Concentration was measured with NanoDrop 2000 Spectrophotometer (Thermo Fisher Scientific). Prior to library preparation, to provide an internal performance control for further steps, 1.75 μg of RNA was mixed with 3.51 μl of 1:99 diluted ERCC RNA Spike-In Control Mix 1 (Ambion). Subsequently, rRNA was depleted using Ribo-Zero Gold rRNA Removal Kit (Human/Mouse/Rat, Illumina) according to the manufacturer’s protocol. Fragmentation and first strand cDNA synthesis were performed as in TruSeq RNA Library Prep kit v2 protocol (Illumina, RS-122-2001, instruction number 15026495 Rev. D), using SuperScript III Reverse Transcriptase (Thermo Fisher Scientific). For second strand synthesis, reaction mixtures were supplemented with 1 μl of 5x First Strand Synthesis Buffer, 15 μl 5x Second Strand Synthesis Buffer (Thermo Fisher Scientific),
0.45 μl 50 mM MgCl, 1 μl 100 mM DTT, 2 μl of 10 mM dUNTP Mix (dATP, dGTP, dCTP, dUTP, 10 mM each, Thermo Fisher Scientific), water to 57 μl, 5 U E. coli DNA Ligase (NEB), 20 U E. coli DNA Polymerase I (NEB), 1 U RNase H (Thermo Fisher Scientific), and incubated at 16°C for 2h. Further steps: purification, end-repair, A-tailing and adapter ligation were performed as described in TruSeq kit protocol with one modification: the first purification eluate was not decanted from the magnetic beads and subsequent steps were performed with the beads in solution. Instead of a new portion of magnetic beads, an equal volume of 20% PEG 8000 in 2.5 M NaCl was added and the DNA bound to the beads already present in the mixture. After the second clean up procedure after adapter ligation, the supernatant was separated from the beads and treated with USER Enzyme (NEB) in 1x UDG Reaction Buffer (NEB) at 37°C for 30 min. The digestion step ensures that the second strand synthetized with dUTP instead of dTTP is removed from cDNA, resulting in strand-specific libraries. The product was amplified using 1 U of Phusion High-Fidelity DNA Polymerase (Thermo Fisher Scientific) in 1x HF Buffer supplemented with 0.2 mM dNTP Mix, and the following primers: PP1 (5’-AATGATACGGCGACCACCGAGATCTACACTCTTTCCCTACACGA-3’), PP2 (5’-CAAGCAG AAGACGGCATACGAGAT-3’). TruSeq kit protocol temperature scheme with 12 amplification cycles and subsequent purification procedure was applied. Enriched library quality was verified using 2100 Bioanalyzer and High Sensitivity DNA kit (Agilent). The libraries’ concentration was estimated by qPCR means with KAPA Universal Library Quantification Kit (Kapa Biosystems), according to the supplied protocol. These libraries were subsequently sequenced using an Illumina HiSeq sequencing platform to the average number of 20 million reads per sample in 100-nt pair-end mode.

Reads were mapped to the mm10 mouse reference genome (GRCm38 primary assembly) using STAR short read aligner with default settings (version STAR_2.5.2) (Dobin, Davis et al., 2013) yielding an average of 62% (polysomal fraction) and 85% (Total RNA sample) of uniquely mapped reads. The quality control, read processing and filtering, visualization of the results, and counting of reads to the Genecode vM6 basic annotation was performed using custom scripts utilizing elements of the HTseq, RSeQC, BEDtools and SAMtools packages (Anders, Pyl et al., 2015, Li, Handsaker et al., 2009, Quinlan & Hall, 2010, Wang, Nie et al., 2016). Differential expression analysis between the NMDA-R stimulated polysome and non-stimulated Total RNA samples was performed using the DESeq2 Bioconductor R package (Love, Huber et al., 2014). The differential expression analysis from the RNA-seq experiment has been correlated with differential MS results.

### RNA isolation from polysomal fractions and qRT-PCR

Before the RNA extraction, external spike-in control mRNA (an *in vitro* transcribed fragment of A. thaliana LSm gene, 10 ng) was added to each of the fractions to control the extraction step. RNA in each fraction was supplemented with linear polyacrylamide (20 μg/ml) and precipitated with 1:10 volume of 3 M sodium acetate, pH 5.2 and 1 volume of isopropanol. After proteinase K digestion RNA was isolated using phenol/chloroform extraction method. Phase separation was performed using Phase Lock Gel Heavy 2 ml Tubes (5Prime). RNA pellets were dissolved in 30 μl of RNase-free water. Equal volume from each RNA sample (6 μl) was reverse transcribed using random primers (GeneON; #S300; 200 ng/RT reaction) and SuperScript IV Reverse Transcriptase (Thermo Fisher Scientific). Next, the abundance of studied mRNAs in different polysomal fractions was analyzed by qRT-PCR using Light Cycler 480 Probes Master Mix (Roche) in a LightCycler480 (Roche). The cDNA was diluted 5× with H2O and 4 μl of each cDNA sample was amplified using a set of custom sequence-specific primers and TaqMan MGB probes in a final reaction volume of 15 μl.

Following TaqMan Gene Expression Assays (Thermo Fisher Scientific) were used: Dnm1l (dynamin 1-like); Suclg1 (succinate-CoA ligase, alpha subunit); Sod2 (superoxide dismutase 2); Ndufa10 (NADH dehydrogenase (ubiquinone) 1 alpha subcomplex 10); Sept4 (septin 4); Agk (acylglycerol kinase); Pdhb (pyruvate dehydrogenase E1, beta subunit); Rab35 (RAB35, member RAS oncogene family); Slc25a18 (solute carrier family 25 member 18); AT3G14080 (U6 snRNA-associated Sm-like protein LSm1).

Relative mRNA levels in different fractions were determined using the ΔCt (where Ct is the threshold cycle) relative quantification method and presented as the % of mRNA in each fraction. Values were normalized to LSm spike-in control.

### Quantification and Statistical Analysis

Unless otherwise noted, statistical analysis was performed using GraphPad Prism 7.0 (GraphPad Software, Inc.). Statistical details of experiments, including the statistical tests used and the value of n, are noted in figure legends.

Proteomics data were statistically analyzed with Scaffold 4 (Proteome Software) platform. Groups were compared with Mann-Whitney test (p<0.05, Benjamin Hochberg correction). Unstimulated synaptoneurosomes were used as a reference sample. Positive fold-change value suggests an increase in the intensity of a given protein as a result of stimulation.

Interactome was prepared and visually edited with CytoScape 3.6.0 (3) platform, using the String application.

For differential analysis of RNA-Seq count data method (Love et al., 2014) has been used. The deseq2 statistical model applies empirical Bayes shrinkage estimation for dispersions and fold change values and Wald test for statistical significance testing. Hits were defined as significantly deregulated when adjusted P values < 0.01.

Statistical analysis of the data from qRT-PCR of polyribosomal fractions was analyzed using one-way ANOVA followed by Sidak’s test for multiple comparisons. The data followed a Gaussian distribution according to the Shapiro-Wilk normality test (p>0.05). Data are presented as means ± SEM.

Data with 2 groups, consisting of pairs of data from independent experiments (high resolution respirometry data) was analyzed using two-tailed paired Student’s t-test. To verify that the difference between pairs follows a Gaussian distribution, D’Agostino & Pearson normality test was used. The effectiveness of pairing was verified by calculating the Pearson correlation coefficient and a corresponding P value. Data are presented as a box-and-whiskers graph.

### Data and Software Availability

The RNA-Sequencing data discussed in this publication have been deposited in NCBI’s Gene Expression Omnibus (Edgar et al., 2002) and are accessible through GEO Series accession number GSE122724 (https://www.ncbi.nlm.nih.gov/geo/query/acc.cgi?acc=GSE122724).

The proteomic data discussed in this publication have been deposited in PRIDE archive px-submissions PXD012746 and PXD012707.

Project accession: PXD012746

Reviewer account details: Username; Password: ld5FdayN

Project accession: PXD012707

Reviewer account details: Username; Password: nkR66pLa

## Acknowledgments

This work was mainly supported by NCN grant Sonata Bis 2014/14/E/NZ3/00375 for MDz; The work in A.C. laboratory was founded by National Science Centre grants OPUS6 2013/11/B/NZ3/00974, OPUS10 2015/19/B/NZ3/03272 and Ministerial funds for science within Ideas Plus program 000263 in 2014-2017.We thank Ben Hur Mussulini for the discussion of experimental procedures.

The equipment used was sponsored in part by the Centre for Preclinical Research and Technology (CePT), a project co-sponsored by European Regional Development Fund and Innovative Economy, The National Cohesion Strategy of Poland.

## Author contributions

MDz, ADz and ACh designed the research. BK, MW, PS, JM, PK performed the research. BK, MW, PS, JM, TMK, PK analyzed the data. BK, DC, PS prepared the figures. DC designed proteomics experiments, run MS-measurements, analyzed data, performed bioinformatics analysis and data interpretation. MDz wrote the paper. BK prepared Star Methods. All authors reviewed and edited the manuscript.

## Conflict of interest

The authors declare that they have no conflict of interest.

---

**Supplementary Figure S1.**
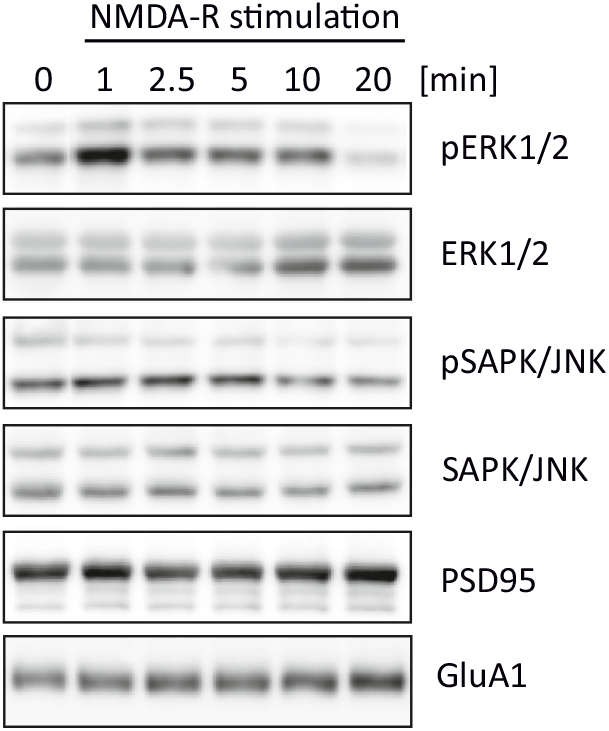
ERK1/2 transient activation in response to NMDAR stimulation in SN. Western blot on synaptoneurosomes verifying activation of selected kinases in response to the stimulation. Chosen NMDA-R stimulation protocol leads to transient phosphorylation of extracellular signal-regulated protein kinases 1 and 2 (ERK1/2). In contrast, stress-activated protein kinase/c-Jun NH2-terminal kinase (SAPK/JNK) are not phosphorylated upon treatment. Antibodies recognizing total ERK1/2 and total SAPK/JNK were used to ensure equal protein levels of analyzed kinases. Anti-PSD95 and anti-GluA1 antibodies were used to verify equal protein loading.

**Supplementary Figure S2.**
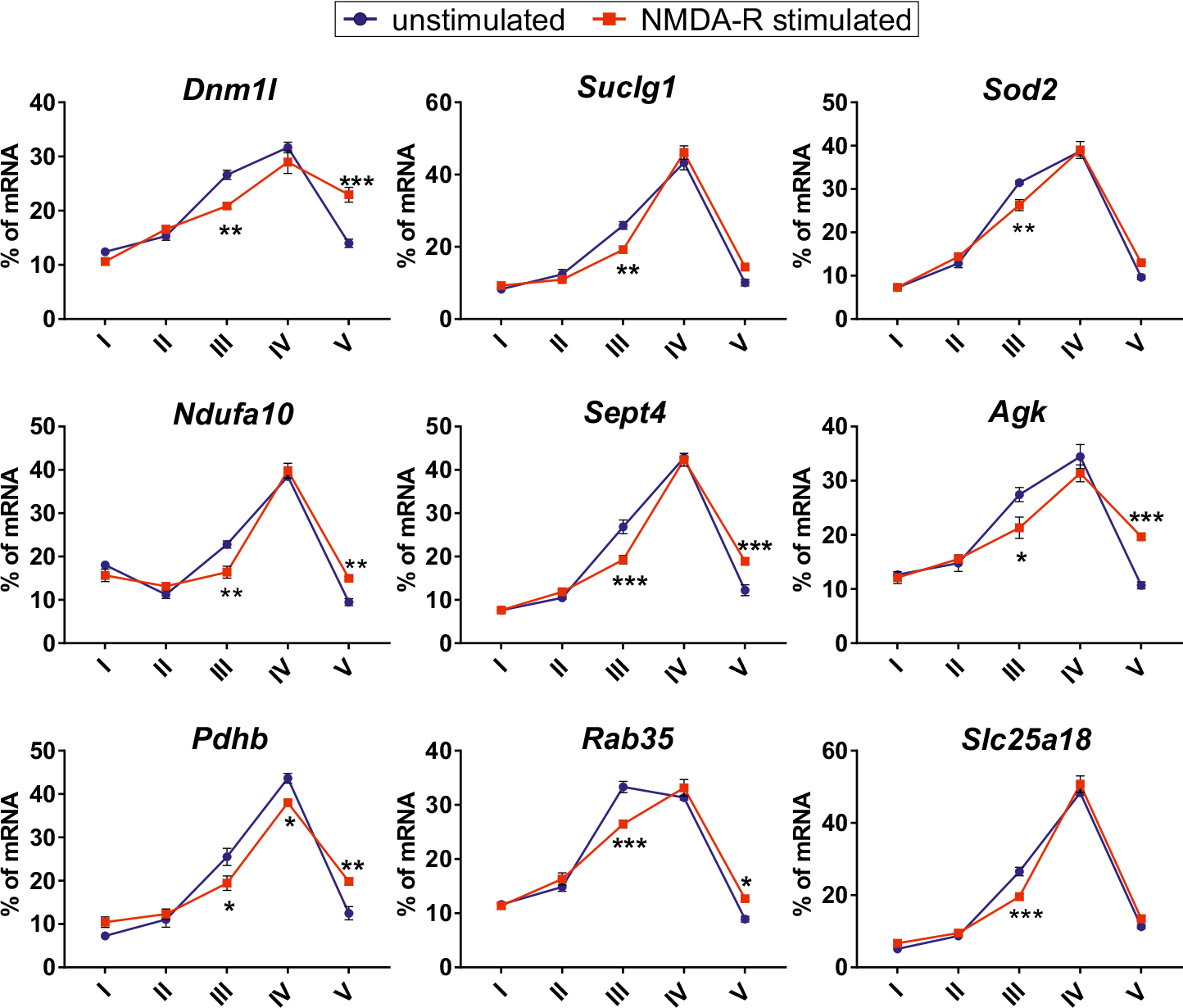
Validation of RNA-seq data using qRT-PCR on polysomal fractions. Graphs show percentage of mRNAs in different fractions from unstimulated and NMDA-R stimulated synaptoneurosomes. Fraction I - monosome, fractions II-III - light polyribosomes and fractions IV-V heavy-polyribosomes. Selected transcripts encoding for mitochondrial proteins shift towards the heavy polyribosomal fractions upon the NMDAR stimulation (n=4; *** p<0.05, ** p<0.01, *** p<0.001; One-way ANOVA, post hoc Sidak’s multiple comparisons test; error bars indicate SEM).

